# Subtle Changes in Clonal Dynamics Underlie the Age-Related Decline in Neurogenesis

**DOI:** 10.1101/206938

**Authors:** Lisa Bast, Filippo Calzolari, Michael Strasser, Jan Hasenauer, Fabian Theis, Jovica Ninkovic, Carsten Marr

## Abstract

Neural stem cells in the adult murine brain have only a limited capacity to self-renew, and the number of neurons they generate drastically declines with age. How cellular dynamics sustain neurogenesis and how alterations with age may result in this decline, are both unresolved issues. Therefore, we clonally traced neural stem cell lineages using confetti reporters in young and middle-aged adult mice. To understand underlying mechanisms, we derived mathematical population models of adult neurogenesis that explain the observed clonal cell type abundances. Models fitting the data best consistently show self renewal of transit amplifying progenitors and rapid neuroblast cell cycle exit. Most importantly, we identified an increase of asymmetric stem cell divisions at the expense of symmetric stem cell differentiation with age. Beyond explaining existing longitudinal population data, our model identifies a particular cellular strategy underlying adult neural stem cell homeostasis that gives insights into the aging of a stem cell compartment.

---

Many adult mammalian somatic tissues are maintained by resident stem and progenitor cell populations and show drastic age-dependent functional decline, which positively correlates with reduced cellular turnover (López-Otín et al., 2013). In mice, the generation of new olfactory bulb (OB) interneurons is sustained by subependymal zone (SEZ) adult neural stem cells (NSCs), whose output substantially decreases during aging (Blackmore et al., 2009; Bouab et al., 2011; Daynac et al., 2016; Mobley et al., 2013; Molofsky et al., 2006; Piccin et al., 2014). Declining neurogenesis has been associated with changes in local or systemic expression of (or responsiveness to) several factors (Chaker et al., 2015; Daynac et al., 2014; Enwere et al., 2004; Katsimpardi et al., 2014; Molofsky et al., 2006; Piccin et al., 2014; Tropepe et al., 1997). Strikingly, it is still unclear if the proliferation of NSCs, and the migration, differentiation and survival of their progeny are affected by age in vivo. The abundance and proliferative activity of NSCs has been reported as decreasing with age (Enwere et al., 2004) or as being mostly unaffected (Daynac et al., 2014, 2016; Shook et al., 2012; Tropepe et al., 1997). Such conflicting views could stem from the different assays employed to evaluate NSC abundance and properties. These comprise *in vitro* assays of cellular behaviors (e.g. neurosphere-forming ability or growth as adherent cultures) known to be significantly affected by exposure to commonly employed mitogens (Costa et al., 2011; Doetsch F, Petreanu L, Caille I, Garcia-Verdugo JM, Alvarez-Buylla A, 2002; Hack et al., 2004) and short-term *ex vivo* analyses of purified cell types (Codega et al., 2014). *In vivo* analyses without clonal lineage tracing (Petreanu and Alvarez-Buylla, 2002) allow for population dynamics snapshots, but are limited in the amount of information they can provide on the progeny of single stem cells.

To overcome these limitations, we recently employed *in vivo* clonal lineage tracing to qualitatively describe the predominant mode of neurogenic NSC activity in the SEZ of adult mice at the age of 2-3 months (from now on called ‘young’ mice) (Calzolari et al., 2015). Our observations support a model of adult OB neurogenesis whereby serial activation of dormant NSCs, followed by a phase of intense neuronal production, is often terminated by NSC exhaustion within a few weeks. While we posited that this process would gradually erode the dormant NSC pool, explaining the age-associated decline in neurogenic activity, it remained unclear if and to which extent changes in proliferation and differentiation during lineage progression play a role.

To tackle this issue, we performed an *in vivo* clonal lineage tracing analysis of adult murine OB neurogenesis at 12-14 months of age (from now on called ‘aged’ mice). At this age neurogenesis is already markedly (3-4 fold) decreased compared to young adult mice, as reflected by the abundance of immature NSC progeny (Bouab et al., 2011; Daynac et al., 2014, 2016; Luo et al., 2006; Molofsky et al., 2006), and overall new OB neuron production (Bouab et al., 2011; Molofsky et al., 2006), thus providing a relevant system. We performed *in vivo* clonal lineage tracing using double hemizygous GLAST^CreERT2^:Confetti transgenic mice (Calzolari et al., 2015; Mori and Tanaka, 2006; Ninkovic et al., 2007), and NSCs were clonally labeled with a single low dose of Tamoxifen (see Supplemental Experimental Procedures). We chose to analyze clones 21 or 56 days post-labeling (dpl; Figure 1A) based on our previous observations in young adult mice (analyzed at 7, 21, 35, and 56 dpl; Figure 1A), which had revealed a clear shift in clonal composition across this time window, from immature clones containing progenitors at earlier timepoints, to clones composed mostly of mature neurons at 56 dpi (Calzolari et al., 2015). We identified clonal components based on a combination of marker expression, localization and cell morphology (Figure 1B) in 46 clones from young mice (reported on in (Calzolari et al., 2015) and 21 clones from aged mice, in total counting 2336 single cells. To our surprise, clone size (Figure 1C) and spatial organization (Figure 1D) did not differ between young and aged mice. Similarly to young animals, clones traced in aged mice showed rapid growth, comprising up to 110 cells already at 21 dpl (Figure 1A). The size, composition and distribution of clusters of TAPs/NBs in the SEZ and proximal rostral migratory stream (RMS) suggested multiple doublings as the basis for lineage amplification (Figure 1E), followed by coherent migration of related NBs (Figures 1F, G and S1A). These observations are similar to the ones in young animals obtained via *in vitro (Costa et al., 2011)* and *in vivo* (*Calzolari et al., 2015*) clonal and population-level analyses (Ponti et al., 2013). Moreover, already at 21 dpl the overall spatial distribution of TAPs, NBs and neurons was compatible with multiple rounds of NSC activation, resulting in the production of bouts of progeny then coherently undergoing maturation and migration (Figure 1G), similar to observations in young animals (Calzolari et al., 2015). Overall clonal maturation dynamics also resembled those observed in young mice; by 21 dpl most clones comprised either only progenitor cells (TAPs/NBs) or progenitors and neurons, with only a minority of clones (2/10) consisting of neurons only (Figure 1A). Eight weeks after labeling (56 dpl) the proportion of clones comprising only neurons had increased (4/11), albeit much less than in young animals, where 7 out of 12 clones consist of only neurons (Figure 1A). These clones were rarely found in association (i.e. in the same hemisphere) with a radial astrocyte (Figure 1H), suggestive of NSC exhaustion being the major mechanism of termination of NSC-derived OB neurogenesis, like in the young SEZ (Calzolari et al., 2015). Finally, the inter- and intra-clonal diversity and distribution of mature neurons in the OB also indicated consistency with the principles deduced from observations in young animals (Calzolari et al., 2015; Fuentealba et al., 2015; Merkle et al., 2013), with mostly subtype-restricted clonal neurogenic activity (Figure 1I,J; Figure S1B-F). These observations revealed that individual NSC clones active in the aged SEZ show no signs of grossly impaired neurogenic activity. This raised the possibility that subtle changes in clonal dynamics may underlie the known decline in overall neurogenic output from the aged SEZ. In order to quantify such features and compare competing hypotheses of clonal dynamics, we mathematically modelled adult neurogenesis at young and middle ages with a stochastic population model (see Supplemental Experimental Procedures) using our clonal data and a limited set of published population-level data, as previously done for other systems (Chabab et al., 2016; Flossdorf et al., 2015; Yang et al., 2015).

**Figure 1:**
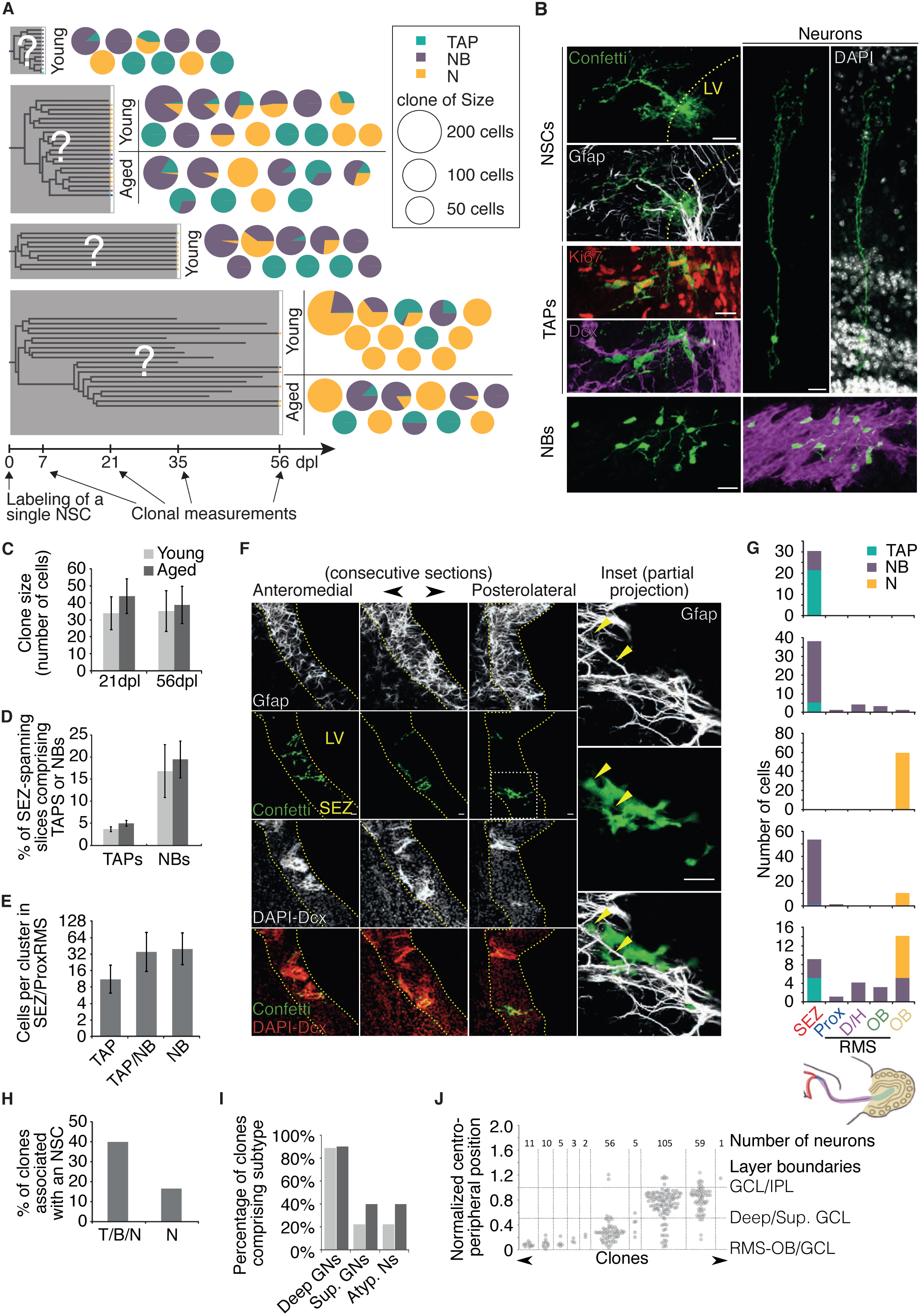
In vivo clonal measurements of neural stem cells (NSCs) in the subependymal zone (SEZ) of young and aged mice. **(A)** Experimental design. The clonal progeny of a single labeled NSC is observed at one of four different time points (7, 21, 35 and 56 days post labeling) in young (white) and aged (grey) mice. The progeny is classified into four cell types: NSC, Transit-amplifying progenitor (TAP), neuroblast (NB) and neuron (N). Pie charts detail the number and composition of clones observed at each time point. Size of pie charts reflect the clone size. **(B)** Examples of cells at distinct stages of neurogenic lineage progression, as labelled in Glast^CreERt2^-Confetti mice. Markers were used to positively identify cell states via GFAP (NSCs) and Dcx (NBs) expression. TAPs and Neurons were defined by a combination of lack of marker expression, localization and morphology. The proliferation marker Ki67 is shown to confirm the TAP identity of SEZ-localized Dcx-negative cells, but was not regularly used to identify cells. Dashed line highlights the LV border. Scale bars 20 μm. **(C)** Average clone sizes at 21 and 56 days post labeling (dpl), for young and aged mice. Data for young mice represent re-plotting of data from Calzolari et al. 2015. We show mean +/-S.E.M. (n=14,12 in young and n=12,11 in aged mice respectively). **(D)** Clonal average percentage of SEZ-encompassing sagittal sections comprising TAPs or NBs, revealing broader distribution for NBs than TAPs, a feature not affected by age. Data for young mice represent re-plotting of data from Calzolari et al. 2015. Error bars represent S.E.M. **(E)** Average size of cell clusters of the indicated compositions, as found in the SEZ/Proximal RMS of aged mice. Error bars represent S.E.M. **(F)** Example of subclonal expansion, showing clone components (Confetti reporter, green) distributed across three consecutive SEZ sections in a 1yo brain. Insets to the right focus on the most posterolateral section, where a single GFAP-positive cell is surrounded by clonally related cells (max-intensity projection of a reduced number of optical sections, to better highlight Confetti/GFAP colocalization). Yellow arrowheads point to GFAP signal in the soma and radial process. Dashed curves indicate SEZ borders, dashed box highlights the inset. LV, lateral ventricle. Scale bars 20 μm. **(G)** Five exemplary clones (aged) showing numbers of cells (y axes) per cell stage (color code) along the SEZ-to-OB axis, based on binning as indicated in the scheme above the panel, depicting a partial sagittal mouse brain section. RMS is subdivided in proximal (Prox), Descending/horizontal limbs (D/H) and RMS-OB (OB). Ocra OB refers to OB locations external to the RMS-OB. **(H)** Percentage of clones, either comprising both progenitors and neurons (T/B/N) or only neurons (N), for which a radial astrocyte sharing the clones Confetti label could be found in the ipsilateral SEZ. **(I)** Percentage of clones comprising the indicated OB neuronal subtypes, for both young and aged mice. Data for young mice are from Calzolari et al. 2015. **(J)** Normalized position of all neurons found in aged mice, subdivided by clone, with number of neurons per clone indicated above the graph.

Murine NSC heterogeneity (besides regionalization (Merkle et al., 2007, 2013)) is well appreciated molecularly and functionally (Codega et al., 2014; Dulken et al., 2017; Llorens-Bobadilla et al., 2015) and evidence exists for interconversion between actively proliferating and temporarily quiescent states (Basak et al., 2012; Costa et al., 2011; Giachino et al., 2014). Dormancy is a recognized feature of the majority of (young adult) NSCs (Shook et al., 2012; Urbán et al., 2016), possibly since late prenatal times (Falk et al., 2017; Fuentealba et al., 2015; Furutachi et al., 2015). We thus modeled the adult neurogenic lineage as comprising three NSC states (fully “dormant” (dS), “quiescent” (qS) and proliferating, “active” (aS) cells), TAPs, proliferating (NB I) and non-proliferating (NB II) neuroblasts and neurons (N) (see Figure 2A, S2A and Supplemental Experimental Procedures for details on model construction). By defining activation and inactivation, proliferation, migration and death rates as parameters for each state, we set up stochastic reaction rate equations that model clonal dynamics. For the three proliferating states (aS, TAP, NB, see Figure 2A), we each allow for four different division strategies: asymmetric (A), symmetric (S), constrained (C), where the proportion of symmetric and asymmetric divisions is regulated by a single parameter p_d_, and unconstrained (U), where any combination of asymmetric division, self-renewal and symmetric differentiation probabilities is allowed, giving rise to an additional parameter (see Figure 2B and Supplemental Experimental Procedures for details). We here define an asymmetric division as a cell division followed by transition of only one daughter cell to the next stage in our model (Figure 2A) before it possibly divides again. Of note, the model does not differentiate whether this transition is coupled to the division, or if it happens some time after the cell has divided. The four different strategies result in 4^3^ = 64 different possible models with varying number of parameters and complexity (Figure 2B). Unknown model parameters were estimated for each model separately by fitting means and variances of modeled TAPs, NBs, and Ns to means and variances of the measured clonal compositions (Figure S2B) using maximum likelihood estimation (Buchholz et al., 2013; Kazeroonian et al., 2016; Stapor et al., 2017). Parameter boundaries and constraints were carefully chosen in accordance with prior knowledge (see Supplemental Experimental Procedures, Table T2). The 64 different models were compared according to the Bayesian Information Criterion (BIC), a score that ranks models based on both their complexity and their ability to explain the observed data (Figure S2C). The ten best performing models (Figure 2C) indicate that changes in the division strategy of active NSCs are required to explain the observed clonal dynamics: While in young mice the best models require symmetric stem cell divisions (allowed by symmetric (S), constrained (C) and unconstrained (U) division strategies), asymmetric (A) stem cell divisions suffice to explain clonal measurements in aged mice (Figure 2C) in the top seven models.

**Figure 2:**
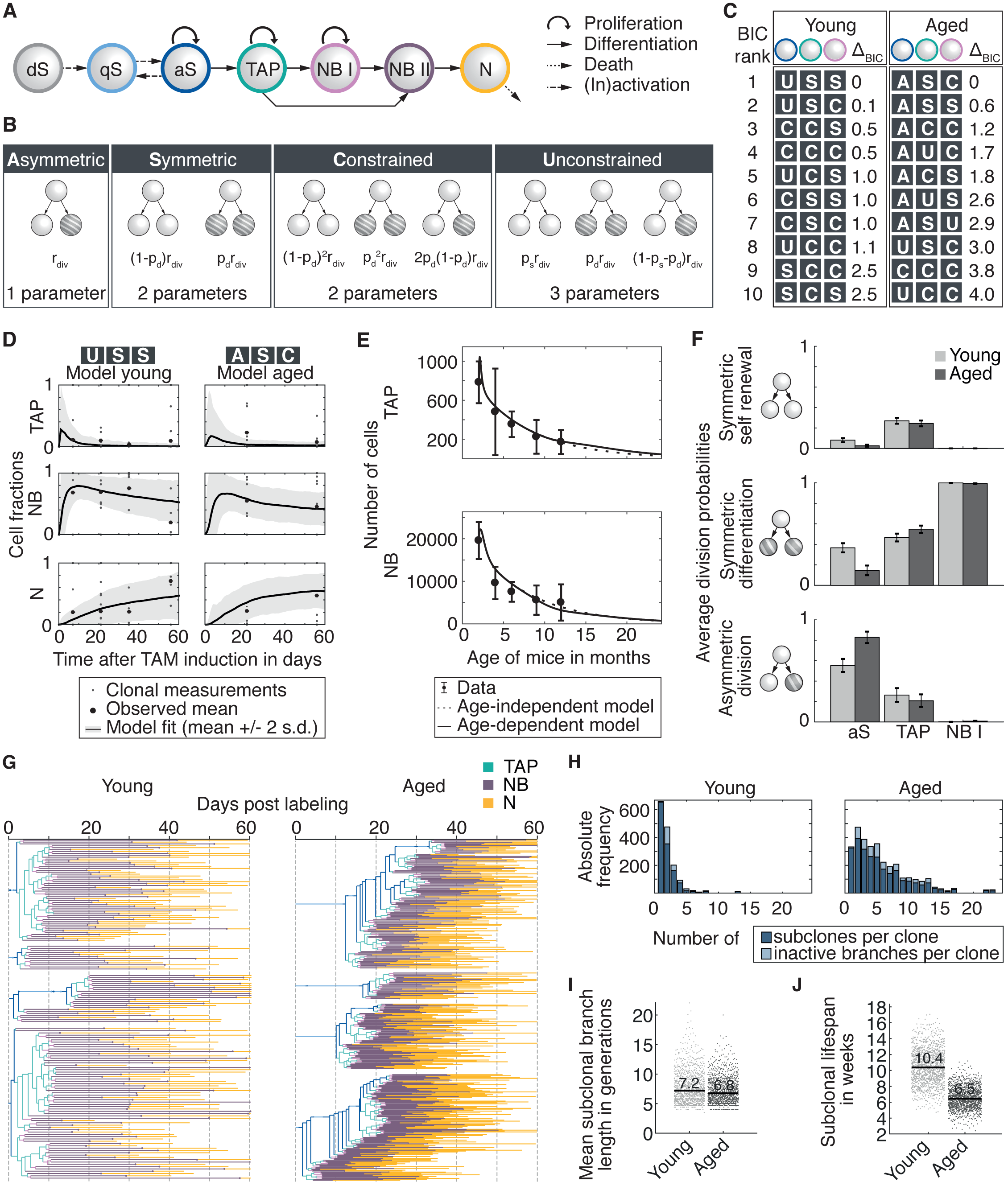
A population model fits the clonal data and predicts increased asymmetric stem cell divisions in aged mice. **(A)**Adult neurogenesis model: The pool of dormant stem cells (dS) is depleted over time. Cells can then be activated and inactivated by switching between the quiescent (qS) and active (aS) state. aS, Transit amplifying progenitors (TAPS) and neuroblasts of type I (NB I) divide. Neuroblasts of type II (NB II) no longer divide and migrate along the SEZ to eventually become neurons (N) that are depleted via cell death. **(B)**Division strategies for dividing cell types: Asymmetric divisions (A) give rise to a daughter cell of the same type and a daughter cell of the subse-quent type, symmetric divisions (S) produce two daughters of the same cell type, constrained divisions (C) assume independent differentiation between sisters, while the unconstrained division (U) is the most flexible strategy. The number of model parameters increases from left to right with equal model complexity for strategies S and C. **(C)** 64 different models are fitted separately to data from young and aged mice and compared via the Bayesian information criterion (BIC). Columns belong to cell types shown in A. Asymmetric stem cell divisions are prevalent in the best 10 models for aged mice. **(D)** Mean cell fractions of best models (solid lines) vs. observed cell fractions (small grey dots) and their mean (large black dots) for TAPs, neuroblasts, and neurons. Model stochasticity is calculated from SSA simulations (gray shaded area, ±2 s.d. errors). **(E)**Predicted cell numbers of age-dependent and age-independent weighted average models (solid and dashed lines) fit to population data from Daynac et al. (mean±2s.d., n>=4 per time point). Data points reflect number of cells per brain hemisphere. Initial conditions are set to earliest observed measurement of the respective cell type. Models include halfway migration of neuroblasts in order to be consistent with the population study data. Data points reflect number of cells per brain hemisphere. **(F)** Division probabilities calculated from all 64 models as a weighted average according to their BIC weights for young (white) and aged (shaded) mice shows strong TAP self renewal (top), rapid NB differentiation (middle), and increased asymmetric stem cell divisions in aged mice (bottom). Error bars indicate ± standard error of the weighted mean (S.E.M._w_). **(G)** n=4 trees simulated according to average models and parameter distributions introduced in (G) for young (left) and aged (right), respectively. **(H)** Absolute frequency of subclones and inactive branches (resulting from aS return to quiescence without any division) per clone in young and aged. Numbers were estimated from n=1000 simulated lineage trees according to the average young and aged model for 100 days. **(I)** Mean subclonal branch length and **(J)** Subclonal lifespan from 1000 simulated lineage trees from the average young and aged model. Medians are shown as a black line.

However, comparing the best-ranking models (see Figure 2D for cell fractions fitted with the best model for young and aged mice) we do not find a single best model; instead, for the top ten models for clones at 2 and 14 months, differences in the BIC values below 4 are observed (Figures 2C and S2C), which are not considered decisive (Kass and Raftery, 1995). Therefore, we derived average models for young and aged mice by weighting resulting parameter estimates with the posterior probability of their model (BIC weight, see Supplemental Experimental Procedures and Figure S2D) to yield robust model predictions. To evaluate the average model for young mice, we compared it to independent population-level data (Daynac et al., 2016; Shook et al., 2012) on the temporal evolution of cell type abundances during aging (Figure 2E). An adaptation of the average parameters from young to aged mice (using Hill kinetics, see Supplemental Experimental Procedures and Figure S2E) leads to an age-dependent model that describes the decrease of TAPs and neuroblasts similarly well and allows for a prediction of cell numbers also for mice beyond 14 months.

Based on the weighted average proportion of symmetric self renewal, symmetric differentiation, and asymmetric division (Figure 2F), we find that asymmetric stem cell divisions are more prevalent in aged compared to young mice (82,8±11,7% vs. 55,3±8,3% in young mice, mean±variance of averaged probabilities), while symmetric differentiation decreases from 36,6±4,1% in young mice to 14,6±8,9% in aged mice. This result still holds if we only consider the data points at 21 and 56 dpl for young mice (Figure S2F). Our models also identify a high, age-independent percentage of symmetric self renewal of TAPs (27,0±1,4% in young, 24,4±1,5% in aged mice), and rapid differentiation of neuroblasts, two properties that are consistent with the existing knowledge about these cell types (Calzolari et al., 2015; Costa et al., 2011; Ponti et al., 2013), while other parameters are estimated to be similar (Figure S2G).

To investigate why the neuronal output diminishes during ageing, despite active stem cells dividing more often asymmetrically, we generated clonal genealogies from the models (see Supplemental Experimental Procedures). We simulated 1000 clones for young and aged mice (Figure 2G) using the inferred weighted average parameters (Figure 2F and S2G-H) and calculated genealogical metrics (Figure S2I) to compare clonal dynamics during aging. Interestingly, increased asymmetric NSC divisions change clonal dynamics in aged mice (Figure 2H-J). Our model thus predicts an increase in asymmetric clonal dynamics resulting in the generation of more, but smaller and shorter-lived subclones in aged mice. While enabling persistent neurogenesis, this produces a reduced neurogenic output at the population level. It is tempting to speculate that the inferred potentially “pro-neurogenic” changes in lineage transition parameters (e.g. increased aS asymmetric division probability) may reflect the action of mechanisms at play to compensate for age-associated neurogenesis-depleting processes, such as the progressive activation and loss of dormant NSCs.

In conclusion, we have performed the first *in vivo* clonal analysis of neural stem cell behavior in aged adult mammals and mathematically modelled adult neurogenesis to define quantitative aspects of lineage transition in young and aged mice. Our model fits the observed data and unveils changes in a restricted set of key parameters. These parameters lead to relatively minor alterations in clonal dynamics, which however explain the observed stronger tendency of young animals to produce mature clones (Figure 1A) and are in agreement with drastic population-level age-related decline in adult OB neurogenesis.

## ACKNOWLEDGEMENTS

We thank M. Goetz (LMU Munich and Helmholtz Zentrum München) for comments on the manuscript, A. Kazeroonian (TUM Munich) and C. Loos (Helmholtz Zentrum München) for computational support.

LB was supported by the German Research Foundation (DFG) within the Collaborative Research Centre 1243, Subproject A17. FC was supported in part by NEURON-ERANET (01EW1604) and DFG (CRC1080) grants to Dr. Benedikt Berninger (UMC Mainz) and by intramural funds to FC.

## AUTHOR CONTRIBUTIONS

FC generated data. LB and FC analyzed data. LB constructed model with FC. LB performed parameter inference and model simulations with support from MS and advice from JH and FT. FC and JN initiated the project with FT. JN and CM supervised the study. LB, FC, JN and CM wrote the paper.

## SUPPLEMENTAL INFORMATION

Supplemental information on our experimental and computational strategy can be found in the SUPPLEMENTAL EXPERIMENTAL PROCEDURES. Code is available at https://github.com/QSCD/NeurogenesisAnalysis

**Figure S1:**
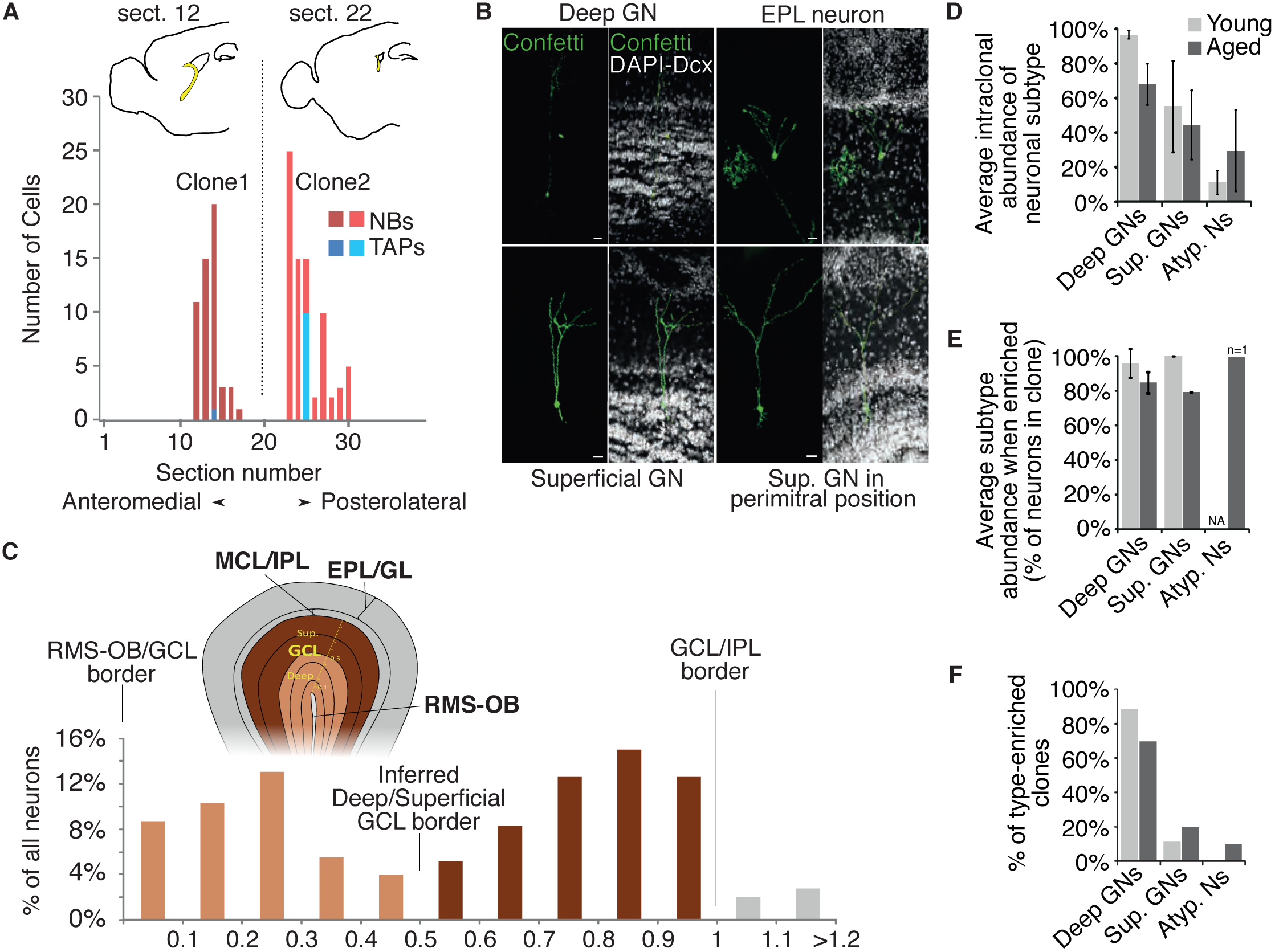
**(A)** Examples of NB/TAP distribution, showing the number and identity of cells, belonging to two distinct clones, found in each sagittal brain slice. Slices are numbered consistently across brains from the medial-most slice. Outlines above the graph refer to the indicated slices and provide an overview of the position and local extent of the SEZ, highlighted in yellow. **(B)** Examples of the variety of OB neuronal subtypes observed. Scale bar 20μm. **(C)** Overall distribution of all OB neurons observed in aged mice; y-axis reports the percentage of neurons found within a given normalized distance bin (shown on the x-axis; 0=RMS-OB, 1=GCL/IPL boundary). Deep and superficial granule neuron domains are clearly distinguishable as distinct distributions. **(D)** Average intraclonal abundance of each neuronal subtype, in young and aged mice. **(E)** Average intraclonal abundance of each neuronal subtype, when prevalent (i.e. when ≥50% of all neurons in the clone), in young and aged mice. **(F)** Percentage of clones being enriched (as defined in E) for the indicated neuronal subtypes, in young and aged mice. Data for young mice are re-plot from Calzolari et al. 2015.

**Figure S2:**
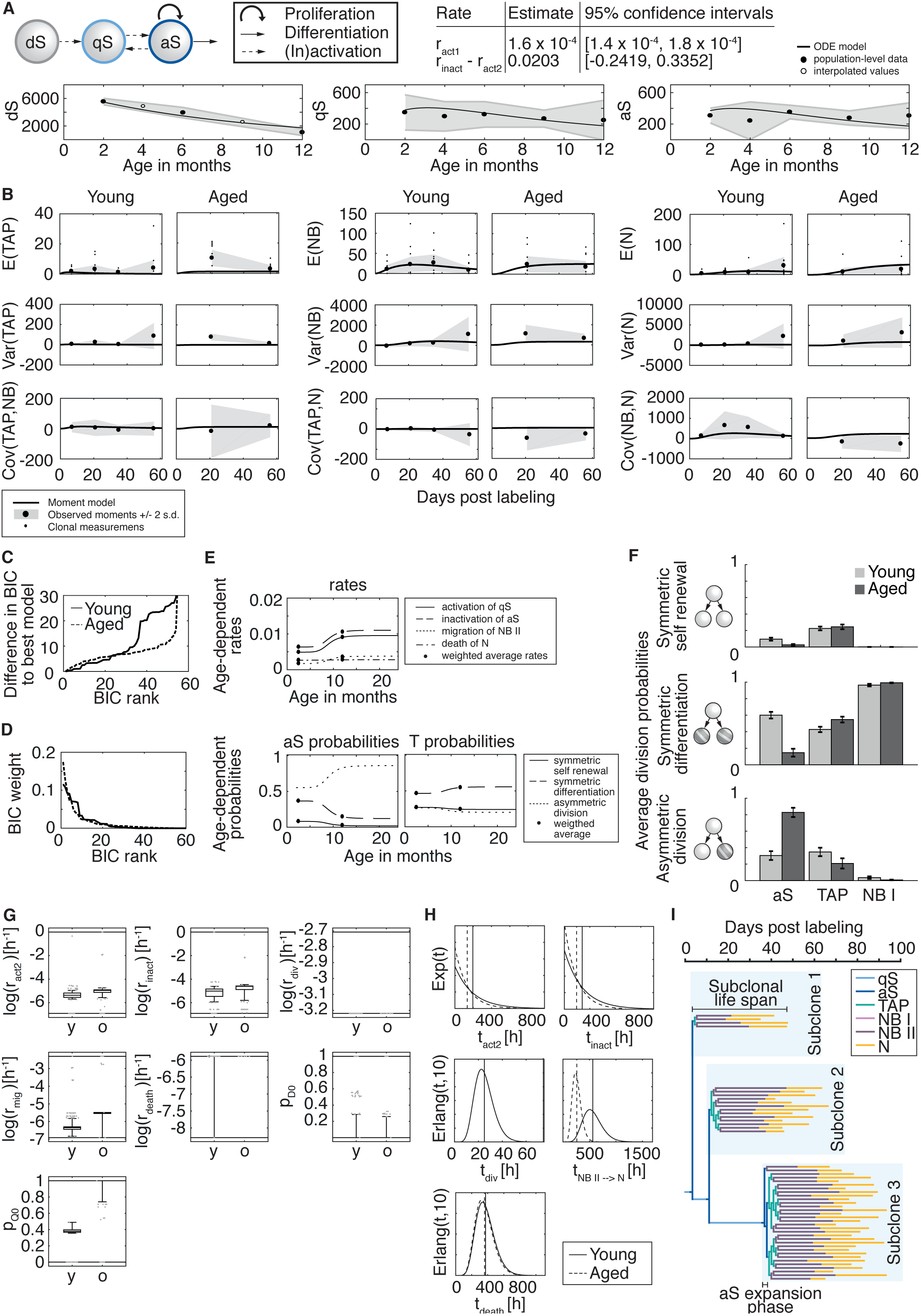
**(A)** Stem cell compartment model (top), fitted to data from Shook et al. (number of active stem cell and total number of stem cells) & Daynac et al. (number of active stem cells and quiescent stem cells), shown as mean ±2 s.d. (n>4 per time point) (middle). Data points reflect number of neural stem cells per brain hemisphere. Rate estimates in table were used to constrain model of neurogenesis. **(B)** Best (rank 1) young and aged models (solid lines) fit to experimental 1st and 2nd order moments (large dots). Standard deviation of experimental moments were calculated via bootstrapping from single clonal observations (small dots in upper row) and shown as ±2 std. error band (grey shaded area). **(C)** Differences in Bayesian Information Criterion (BIC) for all 64 models to the rank 1 model for young (solid line) and aged (dashed line) mice. **(D)** Estimated posterior model probability (BIC weights) for young (solid line) and aged (dashed line) mice indicate that the top ten models dominate. **(E)** Hill function fits to model a smooth age-dependent change in weighted average parameters that differ between young and aged. **(F)** Division probabilities calculated from all 64 models as a weighted average according to their BIC weights for young (white) and aged (shaded) mice using exclusively data of 21 and 56 dpl. Error bars indicate ± standard error of the weighted mean (S.E.M._w_). **(G)** Weighted box plots for log-transformed rates show the weighted probability distribution for parameters resulting from 64 models for young (left) and aged (right). Boxes depict the 1st, 2nd and 3rd quartiles. Outliers correspond to values outside the [2.5%,97,5%]-quantile range (whiskers) and are shown as grey dots. Horizontal lines at top and bottom represent parameter boundaries, which were carefully chosen according to biological plausibility. **(H)** Biologically plausible distributions for (in)activation, division, migration and death times, which are used for tree simulations. Horizontal lines show the mean of distribution (weighted average rates). **(I)** Definition of genealogical metrics.

